# Frataxin deficiency drives cardiac dysfunction and transcriptional dysregulation in Friedreich ataxia iPSC model

**DOI:** 10.1101/2025.08.20.671405

**Authors:** Jarmon G. Lees, Haoxiang Zhang, Lebei Jiao, Anne M Kong, Ren Jie Phang, Li Li, Nan Su, Anthony S. Mukhtar, Alice Pébay, Mirella Dottori, Louise Corben, Martin Delatycki, Roger Peverill, Stephen Wilcox, Jarny Choi, Jeffrey M. Pullin, Davis McCarthy, Jill S. Napierala, Marek Napierala, Shiang Y. Lim

## Abstract

Friedreich ataxia (FRDA) is a progressive neuromuscular degenerative disorder caused by GAA repeat expansions in the *FXN* gene, leading to frataxin deficiency and multisystem pathology. Cardiomyopathy is the leading cause of mortality in individuals with FRDA. To investigate the cellular and molecular mechanisms underlying FRDA-associated cardiac dysfunction, we employed induced pluripotent stem cell (iPSC) lines derived from three individuals with FRDA, each paired with an isogenic control line generated through CRISPR/Cas9-mediated excision of the pathogenic GAA repeat expansion. Correction of the mutation restored *FXN* expression to levels comparable to healthy donor iPSCs, and all lines differentiated efficiently into cardiomyocytes. Functional analysis revealed significant contractile abnormalities in FRDA cardiomyocytes and multicellular cardiac microtissues, including prolonged contraction and relaxation times and faster beating rates, consistent with clinical observations of cardiac contractile dysfunction. FRDA cardiomyocytes also exhibited pathological features such as increased cell size, irregular calcium transients, elevated mitochondrial reactive oxygen species levels, increased mitochondrial fission and increased cell death. These phenotypes were exacerbated by pathological levels of iron supplementation in culture media, highlighting the heightened sensitivity of frataxin-deficient cardiomyocytes to iron-induced metabolic stress. RNA sequencing revealed a distinct transcriptional profile associated with frataxin deficiency. *MEG3* and *PCDHGA10* were consistently dysregulated across all three FRDA-iPSC lines and may represent early molecular markers of FRDA cardiomyopathy. Together, these findings establish a robust human iPSC model of FRDA cardiomyopathy that captures early disease phenotypes and reveals novel molecular targets. This preclinical human model provides valuable insight into the pathogenesis of FRDA and provides a platform for developing early-stage therapeutic interventions.

## Introduction

Friedreich ataxia (FRDA) is one of the most common autosomal recessive ataxias, characterized by progressive neurodegeneration and multisystem involvement, primarily affecting the nervous system and the heart (Keita et al., 2022). The estimated global prevalence of FRDA ranges from 1 in 20,000 to 250,000 individuals depending on geography, with symptom onset typically occurring between 8 and 15 years of age (Reetz et al., 2025). Neurological manifestations commonly include progressive limb and gait ataxia, dysarthria, impaired vibration sense and proprioception, as well as varying degrees of visual and auditory impairment (Keita et al., 2022). Non-neurological features frequently observed in FRDA include scoliosis, diabetes mellitus, and cardiomyopathy. The average life expectancy is approximately 35–40 years, with cardiac complications, particularly arrhythmias and congestive heart failure, being the leading causes of mortality (Indelicato et al., 2024). Currently, there are no approved therapies specifically targeting FRDA-associated cardiomyopathy (Naghipour et al., 2025).

FRDA is caused by mutations in the *FXN* gene located on chromosome 9, which encodes frataxin, a mitochondrial protein essential for cellular energy metabolism (Campuzano et al., 1996). Approximately 96% of affected individuals are homozygous for a large intronic expansion of guanine-adenine-adenine (GAA) trinucleotide repeats within intron 1 of the *FXN* gene (Campuzano et al., 1996; Li et al., 2015). In healthy individuals, the GAA repeat count is typically fewer than 30 per allele, whereas pathogenic expansions exceed 66 repeats, with most individuals with FRDA harboring between 600 and 900 repeats (Reetz et al., 2025). This expansion induces the formation of repressive heterochromatin, marked by increased histone methylation and reduced histone acetylation, which impairs transcriptional initiation and elongation. Consequently, *FXN* mRNA and FXN protein levels are significantly reduced (Chutake et al., 2014).

Frataxin plays a critical role in mitochondrial function, particularly in the biosynthesis of iron-sulfur clusters, which are essential cofactors for multiple mitochondrial enzymes involved in oxidative phosphorylation and cellular energy production (Fox et al., 2019).

Deficiency of frataxin impairs the biogenesis of iron-sulfur clusters and disrupts mitochondrial iron homeostasis, leading to mitochondrial dysfunction and increased oxidative stress. These combined effects result in progressive cellular damage and death in both the nervous and cardiovascular systems (Hanson et al., 2019; Lees et al., 2022).

Patient-derived induced pluripotent stem cells (iPSCs) carrying the pathogenic GAA expansion provide a valuable model for studying FRDA. These cells retain the genetic background and mutation of the donor and can be differentiated into disease-relevant cell types, including cardiomyocytes, enabling the investigation of disease initiation, progression, and therapeutic response in human cellular contexts (Shan et al., 2014). iPSC-based models offer high genetic fidelity and scalability, supporting both mechanistic studies and high-throughput drug screening. However, comparisons with iPSCs from unrelated healthy donors as controls can introduce variability due to differences in genetic backgrounds. Genome editing tools such as CRISPR/Cas9 enable the generation of isogenic control lines by precisely correcting the pathogenic mutation in patient-derived iPSCs, thereby providing a robust platform for controlled experimental studies and direct comparisons between genetically matched cell lines.

In this study, we aimed to characterize the cellular and molecular phenotype of FRDA cardiomyocytes and to identify early markers and mechanisms of FRDA-associated cardiac dysfunction. To this end, we utilized iPSC lines derived from three individuals with FRDA and their corresponding CRISPR/Cas9-corrected isogenic controls. By comparing these paired lines, our objective was to determine how frataxin deficiency resulting from *FXN* GAA repeat expansions affects cardiomyocyte structure, function, and gene expression, and to uncover molecular features that may serve as early indicators or therapeutic targets for FRDA cardiomyopathy.

## Materials and methods

### Human iPSC lines

Fibroblasts from three genetically confirmed adults with FRDA were reprogrammed into iPSC lines (FA1-3), and corresponding CRISPR/Cas9-corrected isogenic control lines (FA1ic-3ic) were generated as previously described (Li et al., 2022) (Table 1). Protocol for derivation of human primary fibroblasts was approved by Institutional Review Board (IRB) at the University of Alabama at Birmingham (IRB N131204003). Both mutant and isogenic iPSC lines were maintained on vitronectin (Thermo Fisher Scientific, MA, USA) coated plates and cultured in StemFlex medium (Thermo Fisher Scientific). A healthy donor iPSC line, CERA007c6 (Hernández et al., 2016), was maintained on vitronectin-coated plates and cultured in TeSR-E8 medium (STEMCELL Technologies, Vancouver, Canada). All human iPSC lines were maintained at 37°C in a humidified atmosphere with 5% CO_2_. Culture medium was refreshed every two to three days, and cells were passaged weekly using ReLeSR (STEMCELL Technologies).

**Table 1.** FRDA patient and iPSC line information. Fibroblast samples from three confirmed individuals with FRDA were reprogrammed into iPSC lines (FA1-3). The disease-causing GAA repeat expansions were corrected using CRISPR/cas9, generating isogenic control iPSC lines (FA1ic-3ic). At the time of biopsy, Participant 1 exhibited hypertrophic cardiomyopathy; Participant 2 showed a severe cardiac phenotype including reduced ejection fraction, asymmetric septal hypertrophy, and arrhythmia; Participant 3 displayed no detectable cardiac abnormalities.

| Individual with FRDA | Sex | GAA1/GAA2 in patient fibroblasts | Derived iPSC lines | Disease state | GAA1/GAA2 in iPSC lines |
| --- | --- | --- | --- | --- | --- |
| 1 | Male | 797/797 | FA1 | FRDA | 867/867 |
|  |  |  | FA1ic | Isogenic control | 0/0 |
| 2 | Male | 404/920 | FA2 | FRDA | 550/830 |
|  |  |  | FA2ic | Isogenic control | 0/0 |
| 3 | Female | 367/780 | FA3 | FRDA | 450/450 |
|  |  |  | FA3ic | Isogenic control | 0/0 |

### Cardiomyocyte differentiation

Cardiomyocytes were derived from iPSCs using previously described methods (Lees et al., 2024; Lyu et al., 2022) with modifications. iPSCs were seeded onto hESC-qualified Matrigel (Corning, NY, USA) coated plates at a density of 2.0×10^5^ cells/cm^2^ in StemFlex medium supplemented with 10 µM Y-27632 (Abcam, Cambridge, UK). After 48 hours, medium was replaced with RPMI 1640 basal medium (Thermo Fisher Scientific) containing B-27 without insulin supplement (Thermo Fisher Scientific), growth factor reduced Matrigel (1:60 dilution; Corning), and 6-8 µM CHIR99021 (STEMCELL Technologies) (designated as day 0). After 24 hours, medium was replaced with RPMI 1640 basal medium containing B-27 without insulin supplement and 5 ng/ml Activin A (Lonza, Basel, Switzerland) for 24 hours. On day 2, the medium was replaced with RPMI 1640 basal medium containing B-27 without insulin supplement and 5 µM IWP2 (Tocris Bioscience, Bristol, UK) for 72 hours. From day 5 onwards, cells were cultured in RPMI 1640 basal medium containing B-27 supplement (Thermo Fisher Scientific) and 20 ng/mL L-ascorbic acid 2-phosphate sesquimagnesium salt hydrate (Sigma-Aldrich, MA, USA) (referred to as CM medium), and the medium was refreshed every 2-3 days. On day 12, cardiomyocytes were dissociated into single cells and split at a 1:4 ratio onto hESC-qualified Matrigel coated plates in DMEM/F-12 GlutaMAX media (Thermo Fisher Scientific) supplemented with 20% fetal bovine serum (Bovogen), 100 µM 2-mercaptoethanol (Thermo Fisher Scientific), 100 µM non-essential amino acids (Thermo Fisher Scientific), 50 U/mL penicillin/streptomycin (Thermo Fisher Scientific) (referred to as CM replating medium), and 10 µM Y-27632. On day 13, the medium was replaced with CM medium. From days 14 to 19, cardiomyocytes were enriched to >95% cardiac troponin T positive cells by culture in glucose-free DMEM medium (Thermo Fisher Scientific) containing 4 mM sodium L-lactate (Sigma-Aldrich). On day 19, cardiomyocytes were dissociated and seeded at a density of 2.5×10^4^ cells/cm^2^ on hESC-qualified Matrigel coated plates in CM replating medium supplemented with 10 µM Y-27632 dihydrochloride. On days 20 and 21, the medium was changed to CM medium, and cardiomyocytes were harvested for endpoints on day 23.

### Endothelial cell differentiation

Human iPSCs were differentiated into CD31 positive endothelial cells according to a previously published protocol (Kong et al., 2019). FA1 and FA1ic iPSCs were dissociated into single cells and seeded onto hESC-qualified Matrigel-coated plates at a density of 1×10^5^ cells/cm^2^ in StemFlex medium supplemented with 10 µM Y-27632. After 24 hours (designated as day 0), the medium was replaced with DMEM/F-12 GlutaMAX medium containing N-2 supplement (Thermo Fisher Scientific), B-27 supplement, 8 µM CHIR99021, and 25 ng/mL BMP4 (STEMCELL Technologies) for 72 hours. On day 3, the medium was replaced with complete StemPro-34 SFM medium (Thermo Fisher Scientific) supplemented with 200 ng/mL VEGF-165 (PeproTech) and 2 µM forskolin (Sigma-Aldrich), and cells were cultured for an additional 72 hours. On day 6, cells were labelled with FITC-conjugated anti-human CD31 antibody (mouse IgG; BD Pharmingen), and CD31 positive cells were isolated via fluorescence-activated cell sorting. Sorted CD31 positive endothelial cells were then expanded on plates coated with 10 µg/mL human fibronectin (Thermo Fisher Scientific) in EGM2-MV medium (Lonza) supplemented with 50 ng/mL VEGF-165 and 10 µM Y-27632 until reaching confluence (designated as passage 0).

### Smooth muscle cell differentiation

Vascular smooth muscle cells were differentiated from human iPSCs according to a published protocol with modifications (Patsch et al., 2015). FA1 and FA1ic iPSCs were dissociated and seeded onto Matrigel-coated plates at a density of 4×10^5^ cells/cm^2^ in StemFlex medium supplemented with 10 μM Y-27632. After 24 hours (designated as day 0), the medium was replaced with N2B27 medium (1:1 ratio mixture of DMEM/F-12 GlutaMAX medium and Neurobasal medium (Thermo Fisher Scientific) plus N-2 supplement and B-27 minus vitamin A supplement (Thermo Fisher Scientific)) supplemented with 8 μM CHIR99021 and 25 ng/mL BMP4. On day 3, the medium was replaced with N2B27 medium supplemented with 10 ng/mL PDGF-BB (PeproTech) and 2 ng/mL Activin A (PeproTech).

On day 4, cells were replated at 8×10^5^ cells/cm^2^ onto collagen (Sigma-Aldrich) coated plates in N2B27 medium supplemented with 2 μg/mL heparin (Sigma-Aldrich) and 2 ng/mL Activin A. From day 4 to day 7, cells were maintained in this medium, after which it was replaced with SmGM-2 medium (Lonza) and continued until day 10.

### Autonomic neuron differentiation

Autonomic neurons were derived from human iPSCs as previously described (Kirino et al., 2018; Lees et al., 2024). Briefly, FA1 and FA1ic iPSCs were seeded into ultra-low attachment round-bottom 96-well plates at a density of 2.0×10^4^ cells/well in E6 medium (Thermo Fisher Scientific) supplemented with 2 µM CHIR99021, 10 µM SB431542 (STEMCELL Technologies), and 10 µM Y-27632 (designated as day 0). On day 3, the medium was replaced by E6 medium supplemented with 20 ng/mL FGF-2 (PeproTech), 50 ng/mL BMP4, 100 nM retinoic acid (Sigma-Aldrich) and 50 U/mL penicillin/streptomycin. The medium was refreshed on day 7. On day 10, neurospheres were collected and dissociated with TrypLE™ Select Enzyme (Thermo Fisher Scientific), then re-seeded into 6-well ultra-low attachment plate (Corning) in Neurobasal Plus basal medium (Thermo Fisher Scientific) supplemented with N-2 Plus supplement (Thermo Fisher Scientific), B-27 Plus supplement (Thermo Fisher Scientific), GlutaMAX (Thermo Fisher Scientific), 20 ng/mL bFGF, 20 ng/mL EGF (PeproTech), 50 ng/mL BMP4, and 2 µg/mL heparin (Sigma-Aldrich). The medium was refreshed on day 14. Beginning on day 18, cultures were maintained in Neurobasal Plus medium supplemented with N-2 Plus supplement, B-27 Plus supplement, 10 ng/mL BDNF (PeproTech), 10 ng/mL GDNF (PeproTech), and 10 ng/mL NGF (PeproTech), with medium changes every 3-4 days to support neuronal maturation. On day 32, neurospheres were dissociated and either cryopreserved or used for the construction of cardiac microtissues.

### Cardiac fibroblast differentiation

Cardiac fibroblasts were differentiated from iPSCs according to a previously published method (Giacomelli et al., 2020; Lees et al., 2024). iPSCs were dissociated into single cells and seeded onto Matrigel-coated plates at a density of 2.5×10^4^ cells/cm^2^ in StemFlex medium supplemented with 10 μM Y-27632. After 24 hours (designated as day 0), the medium was replaced with STEMdiff™ APEL™ 2 medium (STEMCELL Technologies) supplemented with 20 ng/ml BMP4, 20 ng/ml Activin A, and 1.5 µM CHIR99021 for 3 days. On day 3, the medium was replaced with STEMdiff™ APEL™ 2 medium containing 30 ng/ml BMP4, 1 µM retinoic acid, and 5 µM IWP-2. On day 6, the medium was replaced with STEMdiff™ APEL™ 2 medium containing 30 ng/ml BMP4 and 1 µM retinoic acid. On day 9, cells were dissociated and replated at a density of 1.5×10^4^ cells/cm^2^ onto human fibronectin-coated plates in STEMdiff™ APEL™ 2 medium containing 10 μM SB431542 and 10 μM Y-27632. On day 13, epicardial cells were dissociated and replated at 2.5×10^4^ cells/cm^2^ onto 0.1% porcine gelatine-coated plates and cultured in STEMdiff™ APEL™ 2 medium containing 10 ng/ml FGF-2 and 10 μM Y-27632. The medium was changed every 2 days, omitting Y-27632 after initial replating. On day 20, the medium was switched to FGM-3 medium (Lonza) and refreshed every 2-3 days thereafter.

### Cardiac microtissue construction

Engineered multicellular cardiac microtissues were constructed according to previously published protocols with modifications (Lyu et al., 2022; Yin et al., 2025). Briefly, day-19 FA1 or FA1ic cardiomyocytes were seeded onto Matrigel-coated 48-well Nunc^TM^ UpCell^TM^ plates (Thermo Fisher Scientific) at a density of 1.2×10^5^ cells/cm^2^ (equivalent to 2.0×10^5^ cells per cardiac microtissue) in CM replating medium supplemented with 10 μM Y-27632. After 24 hours, FA1 or FA1ic iPSC-derived endothelial cells (5.0×10^4^ cells/cm^2^), smooth muscle cells (5.0×10^3^ cells/cm^2^), autonomic neurons (2.0×10^4^ cells/cm^2^), and cardiac fibroblasts (5.0×10^3^ cells/cm^2^) were seeded onto the cardiomyocyte layer. The co-culture was maintained in cardiac microtissue medium, composed of a 1:1:1:1:1 ratio of CM medium, endothelial EGM2-MV medium, smooth muscle cell SmGM-2 medium, autonomic neuron medium (DMEM/F12 supplemented with 10% FCS), and cardiac fibroblast FGM-3 medium. This medium was further supplemented with 50 ng/mL VEGF-165, 10 ng/mL BDNF, 10 ng/mL GDNF, 10 ng/mL NGF, and 10 μM Y-27632. After 24 hours, the UpCell^TM^ plates were brought to room temperature to release the intact cell sheet, which were then transferred to ultralow attachment plates (Sigma-Aldrich) containing cardiac microtissue medium (without Y-27632) for 24 hours. The resulting spheroids were then embedded in 15 µL of growth factor reduced Matrigel and cultured in cardiac microtissue medium (without Y-27632). Cardiac microtissues were maintained in a humidified CO_2_ incubator on an orbital shaker at 60 rpm, with medium changes every 2-3 days.

### Iron-induced metabolic stress

To simulate iron-induced metabolic stress conditions, 200 μM of iron sulfate was added to the CM medium from day 21 to day 23. Iron sulfate was prepared as a 100 mM stock solution by dissolving 20 mg of iron sulfate tetrahydrate (Sigma-Aldrich) in DPBS containing 0.1% of bovine serum albumin (Sigma Aldrich) adjusted to pH 5.5 (Lee et al., 2014; Lee et al., 2016).

### Reverse transcription-quantitative polymerase chain reaction (RT-qPCR)

Total RNA was extracted from iPSCs and cardiomyocytes using Tri-Reagent (Thermo Fisher Scientific), followed by RNA precipitation with chloroform and isopropanol (Sigma-Aldrich). cDNA was synthesized using a high-capacity cDNA reverse transcription kit (Applied Biosystems, CA, USA). Quantitative PCR was performed in duplicate using TaqMan Universal Master Mix, the QuantStudio 6 Flex Real-Time PCR system, and TaqMan Gene Expression Assays for *FXN* (Hs00175940_m1), *MEG3* (Hs00292028_m1), *PCDHGA10* (Hs00259371_s1), and *GAPDH* (Hs99999905_m1) (Applied Biosystems, CT, USA). Relative mRNA expression was quantified using the comparative CT method (2^-ΔΔCt^) method, normalized to *GAPDH*.

### Frataxin protein expression

Relative frataxin protein expression was determined using the Frataxin Protein Quantity Dipstick Assay Kit (ab109881, Abcam) according to the manufacturer’s protocols. Briefly, 4 µg of total protein from iPSCs or cardiomyocytes, quantified using the Pierce™ BCA Assay (Thermo Fisher Scientific), was applied to microplate wells pre-coated with dried gold-conjugated frataxin antibody and incubated for 5 minutes at room temperature. The samples were then wicked onto dipsticks, allowed to dry at room temperature, and imaged using a flatbed scanner. Band intensities were quantified using ImageJ software.

### Immunofluorescence

Cardiomyocytes were fixed (10% neutral buffered formalin; Trajan Scientific) for 20 minutes at room temperature and permeabilized (0.2% Triton X-100) for 10 minutes. After blocking with serum-free blocking solution (Dako, VIC, Australia) for 10 minutes, cells were stained with the specified primary antibodies overnight at 4°C, followed by secondary antibodies and 1 μg/mL of DAPI (Invitrogen) for 1 hour at room temperature.

#### Cardiomyocyte hypertrophy

To assess cardiomyocyte hypertrophy, fixed cardiomyocytes were immunostained with cardiac troponin-T (2 µg/mL, rabbit polyclonal, ab45932, Abcam) and α-actinin (25 µg/mL, mouse monoclonal, A7811, Sigma-Aldrich). The cell boundaries of cardiomyocytes positive for both cardiac troponin T and α-actinin were manually outlined, and their cell surface area was quantified using Image J software.

#### Mitochondrial morphology

Mitochondrial morphology was assessed by Hsp60 immunostaining of cardiomyocytes cultured on glass coverslips, as previously described (Hoque et al., 2018; Rosdah et al., 2022). Fixed cells were stained with α-actinin and Hsp60 (2 µg/mL, rabbit polyclonal, ab46798, Abcam). Coverslips were mounted onto glass slides using fluorescence mounting medium (Dako), and images were acquired using an Olympus BX61 fluorescence microscope at 600x magnification. For each replicate, images were captured from 10 randomly selected fields of view, and a minimum of 60 α-actinin-positive cardiomyocytes were analyzed.

Mitochondrial morphology was classified as either predominantly networked or fragmented, based on whether more than 50% of the mitochondria in a given cell exhibited that specific morphology. For each condition, the percentages of cells classified as having networked or fragmented mitochondria were calculated, summing to 100%.

#### Multicellular cardiac microtissues

Multicellular cardiac microtissues were fixed in 10% neutral buffered formalin for 2 hours at room temperature, then dehydrated in 20% sucrose solution for 24 hours. Dehydrated samples were embedded in Optimal Cutting temperature compound (Sakura Finetek, CA, USA), and cryosectioned at 10 µm thickness. Sections were permeabilized with 0.2% Triton X-100 and blocked using Protein Block (Dako) before incubation with primary antibodies: cardiac troponin T (2 μg/mL, rabbit polyclonal, Abcam), cardiac troponin T (4 μg/mL, mouse monoclonal, ab8295, Abcam), CD31 (2 μg/mL, mouse monoclonal, M0823, Dako), and peripherin (2 μg/mL, mouse monoclonal, sc377093, Santa Cruz). Following primary antibody incubation, sections were stained with secondary antibodies: Alexa Fluor 488-conjugated goat-anti-rabbit or goat-anti-mouse (10 μg/mL, A11001/A11008, Invitrogen), and Alexa Fluor 594-conjugated goat-anti-mouse or goat-anti-rabbit (10 μg/mL, A11005/A11012, Invitrogen). Nuclei were counterstained with 1 μg/mL of DAPI (Invitrogen). Epifluorescence images were acquired using an Olympus BX61 upright microscope and analySIS software.

### Contraction analysis

Cardiomyocyte and cardiac microtissue contractions were recorded using an inverted Olympus IX-71 microscope (Claridge et al., 2023; Lees et al., 2024). Videos were recorded at either 30 frames per second (for cardiac microtissues) or 60 frames per second (for 2D cardiomyocytes) for 15 seconds or until at least ten complete contraction cycles were captured. Contraction analysis was performed using the MUSCLEMOTION plugin (Sala et al., 2018) in ImageJ software, with the following setting: frames/second (30 or 60), speedWindow (2), default noise reduction, automatic reference frame, and automatic peak fitting. Beat-to-beat variability was quantified by calculating the root mean square of successive differences (RMSSD) from 10 successive beats using the formula: 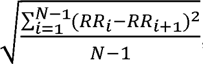, where N is the number of beats, and RR represents the interval between adjacent contraction peaks (Yin et al., 2025).

### Field potential duration

Cardiomyocyte electrophysiological activity was evaluated using a multielectrode array (MEA) recording system (Multichannel Systems, Reutlingen, Germany), as previously described (Lees et al., 2019). Day 19 cardiomyocytes were seeded onto MEA plates pre-coated with hESC-qualified Matrigel and cultured for 48 hours. Extracellular field potentials were recorded for 5 minutes, and data were analyzed offline using MC Rack software (version 4.3.5) to determine RR intervals and extracellular field potential duration (FPD).

FPD values were normalized using Fridericia’s correction formula: corrected FPD = FPD/(RR interval)^1/3^.

### Calcium transient analysis

Calcium transient amplitude and regularity were assessed using the calcium-sensitive indicator Fluo-4 acetoxymethyl ester (Thermo Fisher Scientific). Fluo-4 was prepared at a final concentration of 1 μM in Hanks’ Buffer Saline Solution with Mg^2+^ and Ca^2+^ (HBSS++, Thermo Fisher Scientific). Cardiomyocytes were incubated with Fluo-4 at 37°C in a humidified CO_2_ incubator, protected from light, for 20 minutes. After incubation, cells were washed twice with HBSS++ and imaged immediately in fresh HBSS++ using an inverted Olympus IX-71 microscope. Calcium transients were analyzed by plotting corrected total cell fluorescence of Fluo-4 signal over time. Peak amplitude was quantified as the change in fluorescence intensity relative to baseline (Δ(F-F_0_)) across three consecutive peaks. Calcium traces displaying abnormal spike morphology or irregular rhythmicity were classified as irregular, based on published criteria (Gaballah et al., 2022; Hwang et al., 2020), and expressed as a percentage of total cardiomyocytes analyzed. For each biological replicate, calcium transients were recorded from 20 cardiomyocytes sampled from at least 3 random fields.

### Cell death assay

Cardiomyocyte viability was assessed using propidium iodide and Hoechst 33258 staining. Cells were incubated with 5 μg/mL of propidium iodide and 1 μg/mL of Hoechst 33258 (Sigma-Aldrich), prepared in HBSS++, for 45 minutes at 37°C in a humidified CO_2_ incubator, protected from light. Fluorescent images were captured using an inverted Olympus IX-71 microscope. The number of dead cells (propidium iodide positive) was counted and expressed as a percentage of the total number of cells (Hoechst 33258 positive). At least 100 cells per condition were analyzed in each independent experiment.

### Mitochondrial ROS measurement

Mitochondrial ROS levels in cardiomyocytes were assessed using the MitoSOX™ Red dye (Thermo Fisher Scientific). Cells were incubated with 5 μM MitoSOX™ Red in HBSS++ for 20 minutes at 37°C in a humidified CO_2_ incubator, protected from light. Following incubation, cells were washed once with HBSS++ and imaged immediately in fresh HBSS++ using an inverted Olympus IX-71 microscope at 200x magnification. The corrected total cell fluorescence was quantified for 100 cardiomyocytes from 3 random fields per group in each independent experiment.

### Mitochondrial membrane potential measurement

Mitochondrial membrane potential was assessed using tetramethylrhodamine methyl ester (TMRM, Thermo Fisher Scientific). Cardiomyocytes were incubated with 10 nM TMRM in HBSS++ (non-quenching concentration) for 20 minutes at 37°C in a humidified CO_2_ incubator, protected from light. Imaging was performed immediately after incubation, without a wash step, using an inverted Olympus IX-71 microscope at 200x magnification. The corrected total cell fluorescence was quantified for 100 cardiomyocytes across 3 random fields for each group in each independent experiment.

### RNA sequencing

Total RNA was extracted from cardiomyocytes derived from all three FRDA and isogenic control iPSC lines across three independent experiments using the Cytivia Mini RNA Extraction Kit (Cytivia, MA, USA), following the manufacturer’s instructions. Residual genomic DNA was removed by treatment with TURBO DNase (Thermo Fisher Scientific). RNA integrity was assessed using the Agilent 2100 Bioanalyzer (Agilent Technologies), and all samples exhibited an RNA Integrity Number (RIN) of 10.

Library preparation was performed using 100 ng of input RNA per sample with the TruSeq Stranded Total RNA Library Prep Kit (Illumina, CA, USA) as per manufacturer’s instruction. The library was quantified using the Agilent Tapestation and the Qubit™ RNA assay kit for Qubit 2.0® Fluorometer (Life technologies). Unique dual indexes (Illumina TruSeq RNA UD Indexes) were used to barcode individual libraries. The indexed libraries were then prepared and diluted to 750pM for paired end (2x 101 base) sequencing on a NextSeq2000 instrument using the P3 200 cycle kit using v3 chemistry (Illumina, CA, USA) as per manufacturer’s instructions. The base calling and quality scoring were determined using Real-Time Analysis on board software v2.4.6, while the FASTQ file generation and de-multiplexing utilised bcl2fastq conversion software v2.15.0.4.

RNA sequencing (RNA-seq) data were processed and analyzed using the ARMOR (Automated Reproducible Modular Workflow for Preprocessing and Differential Analysis of RNA-seq Data) pipeline (Orjuela et al., 2019). Following preprocessing steps, including read quality assessment and adapter trimming, reads were aligned to the reference genome and quantified to obtain gene-level expression counts. For differential gene expression analysis, the workflow employed the edgeR statistical package to estimate dispersions and fit generalized linear models to assess differential expression between experimental conditions (Chen et al., 2025). Genes with a false discovery rate (FDR) below a specified threshold (FDR < 0.05) were considered significantly differentially expressed. All analyses were performed with default parameters unless otherwise specified, ensuring consistency and reproducibility across samples.

Pathway enrichment analysis was performed using the PANTHER Overrepresentation Test (release date: 2024-08-07) via the PANTHER online platform (Mi et al., 2019) and visualised using SRplot (Tang et al., 2023). Annotations were based on the GO Ontology database [DOI: 10.5281/zenodo.15066566], released on 2025-03-16. The input gene list included all differentially expressed genes (DEGs) with an FDR < 0.05 and an absolute log fold change > 2. The reference background included all identified DEGs. Enrichment was assessed using Fisher’s Exact Test, and multiple testing correction was applied using the FDR method.

### Statistics

Data are presented as mean ± standard error of the mean (SEM). Statistical significance was evaluated using either an unpaired Student’s *t*-test or a two-way ANOVA, followed by Fisher’s Least Significant Difference test for planned pairwise comparisons, as appropriate. A p-value < 0.05 was considered statistically significant.

## Results

### Restoration of *FXN* mRNA and FXN protein expression in CRISPR/Cas9-corrected isogenic control iPSCs and cardiomyocytes

To model FRDA, iPSC lines were established from three individuals with FRDA (FA1–3), along with isogenic controls (FA1ic–3ic) generated via CRISPR/Cas9-mediated excision of the pathogenic GAA repeat expansion (Table 1). Both FRDA and isogenic control iPSC lines displayed morphology typical of pluripotent stem cells, forming flat, compacted colonies with no discernible signs of spontaneous differentiation under pluripotency-maintaining conditions. Removal of the GAA repeat expansion resulted in a significant increase in FXN mRNA and protein expression in all three isogenic control iPSCs, restoring levels comparable to a healthy donor iPSC line (Figure 1A-B).

**Figure 1.**
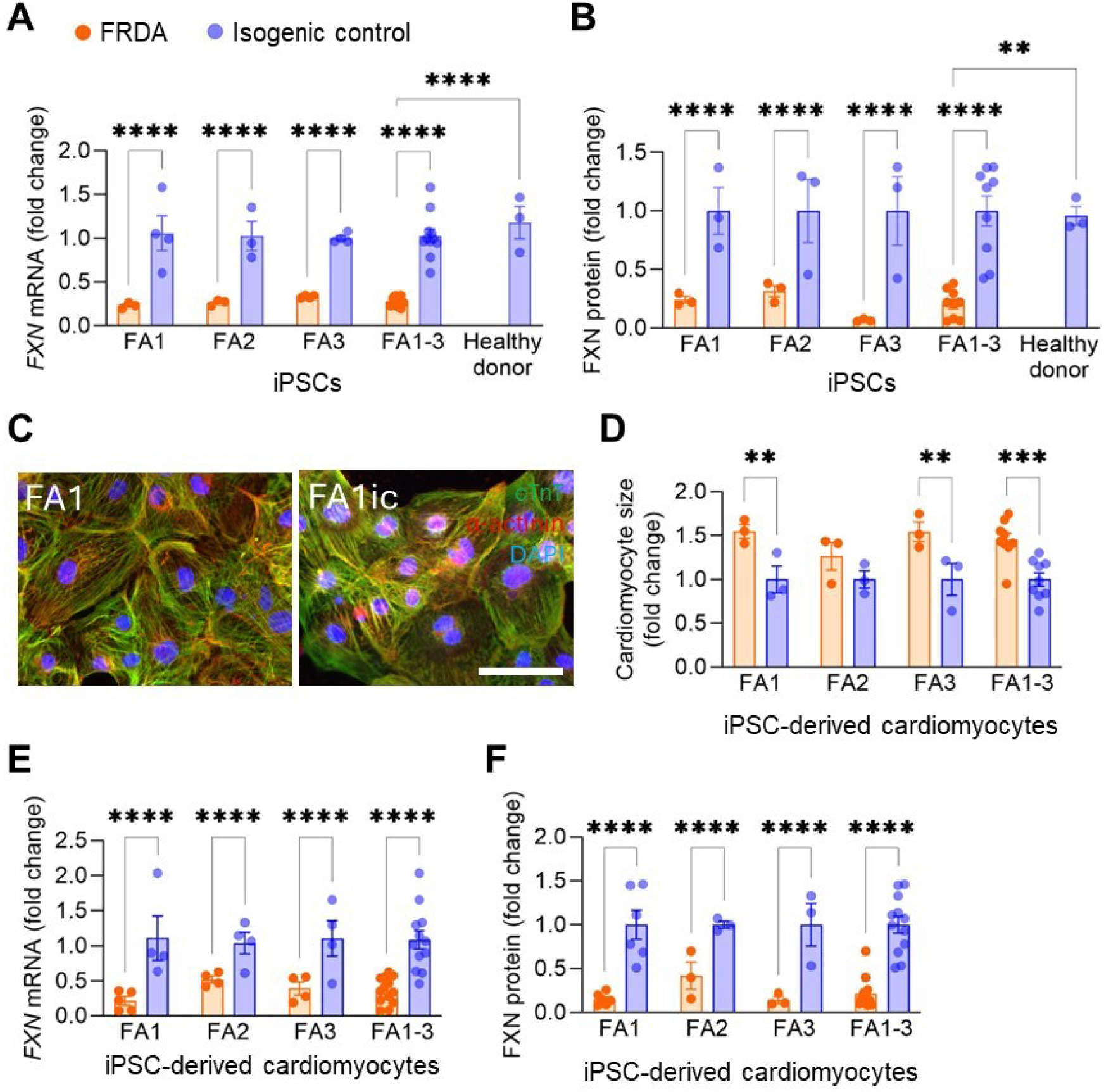
Friedreich ataxia iPSCs and cardiomyocytes exhibit reduced FXN expression compared to isogenic controls. (A-B) FXN mRNA (A) and protein (B) expression levels in FRDA patient-derived iPSC lines, isogenic control iPSC lines, and a healthy donor iPSC line (n = 3-4 independent experiments). (C) Representative immunofluorescent images of iPSC-derived cardiomyocytes stained with cardiac troponin T (green), α-actinin (red), and DAPI (blue). Scale bar = 50 µm. (D-F) Quantification of cardiomyocyte size (D), *FXN* mRNA expression (E), and FXN protein expression (F) in FRDA and isogenic control iPSC-derived cardiomyocytes (n = 3-6 independent experiments). Data are presented as mean ± SEM for individual iPSC lines (FA1, FA2, FA3) and a pooled average across all 3 lines (FA1-3). **p<0.01, ***p<0.001, ****p<0.0001 by unpaired t-test.

Cardiomyocytes were successfully differentiated from both FRDA and isogenic control iPSC lines. Across all lines, cardiomyocytes uniformly expressed cardiac-specific markers, including cardiac troponin T and α-actinin (Figure 1C). A significant increase in cardiomyocyte cell size, suggestive of pathological hypertrophy, was observed in two of the three FRDA lines (Figure 1D). Importantly, FXN mRNA and protein expression levels in the differentiated cardiomyocytes mirrored those observed in the undifferentiated iPSCs, with isogenic control cardiomyocytes showing significantly higher FXN expression compared to their FRDA counterparts (Figure 1E-F).

### Friedreich ataxia iPSC-derived cardiomyocytes exhibit contractile dysfunction

To determine whether FRDA iPSC-derived cardiomyocytes recapitulate the cardiac contractile dysfunction observed in individuals with FRDA, contractile function was assessed by measuring total contraction duration (total time taken to contract and relax), time to peak contraction (a surrogate for systolic function), and relaxation time (a surrogate for diastolic function). Compared to their isogenic control cardiomyocytes, FRDA cardiomyocytes from FA2 and FA3 iPSC lines exhibited significantly prolonged total contraction duration, time to peak, and relaxation time (Figure 2A-D). In contrast, the FA1 cardiomyocytes did not exhibit prolonged total contraction duration or time to peak (Figure 2B-C), although relaxation time was significantly prolonged (Figure 2D). When data from all three FRDA lines (FA1-3) were pooled, a significant overall prolongation of all contractile parameters was observed, consistent with impaired systolic and diastolic function. Beating rate (beats per minute, BPM) was significantly lower in the FA1 and FA3 FRDA lines compared to their isogenic controls, but no difference was observed in the FA2 line (Figure 2E). Beat rate variability, an indicator of arrhythmic risk, did not differ between FRDA and isogenic control cardiomyocytes (Figure 2F).

**Figure 2.**
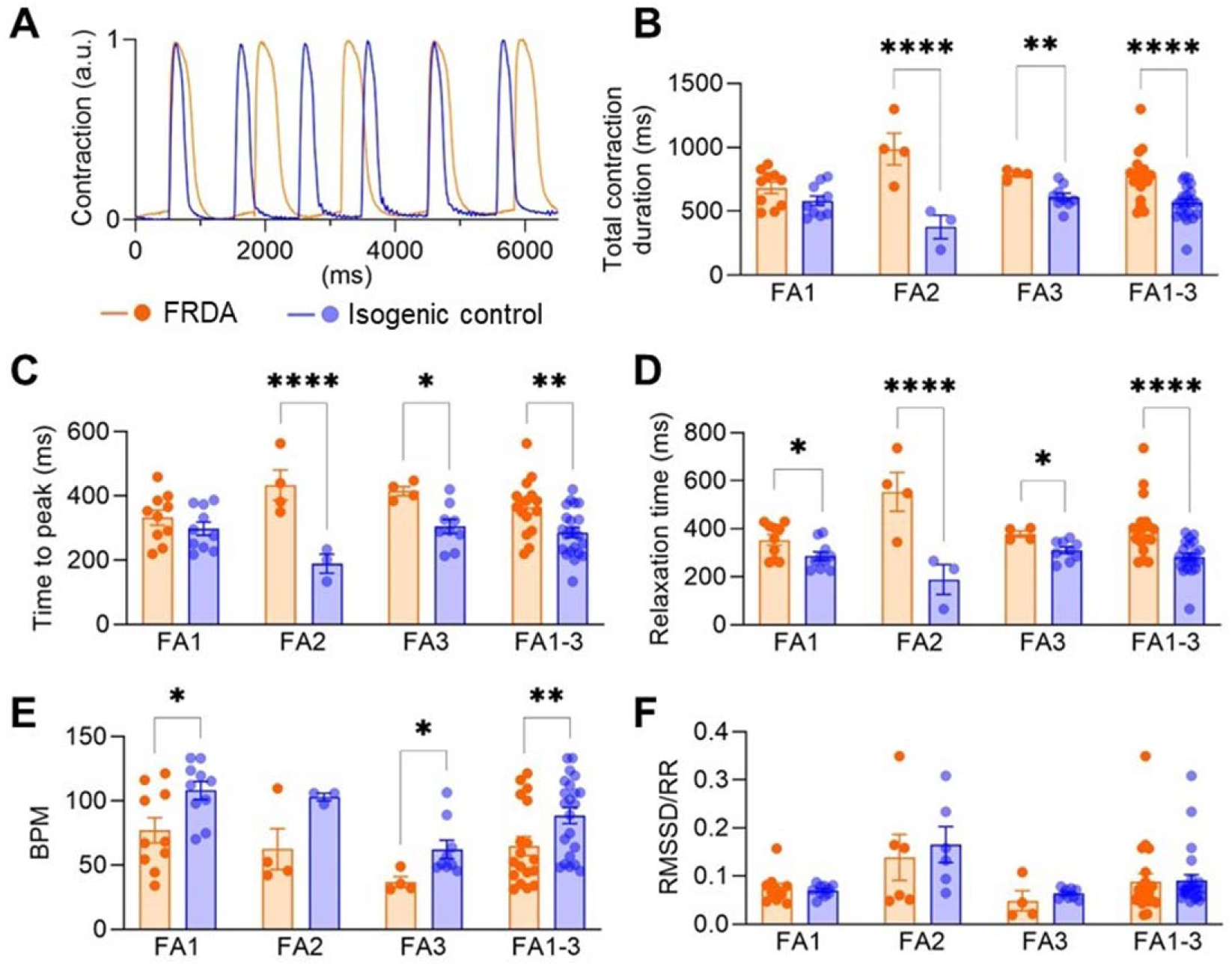
Contractile dysfunction in Friedreich ataxia iPSC-derived cardiomyocytes. (A) Representative contraction trace of FRDA and isogenic control cardiomyocytes (a.u. = arbitrary units). (B-E) Quantification of total contraction duration (B), time to peak contraction (C), relaxation time (D), beating rate (BPM) (E), and beat rate variability (RMSSD/RR) (F) in FRDA and isogenic control cardiomyocytes (n = 3-5 independent experiments). Data represent mean ± SEM for individual iPSC lines (FA1, FA2, FA3) and the pooled average across all 3 lines (FA1-3). *p<0.05, **p<0.01, ****p<0.0001 by unpaired t-test.

### The incidence of irregular cardiomyocyte calcium oscillations is increased in Friedreich ataxia iPSC-derived cardiomyocytes

Calcium signalling plays a critical role in the contraction and relaxation of cardiomyocytes (Landstrom et al., 2017). In individuals with FRDA, abnormalities in calcium signalling, both in intensity and oscillation stability, have been reported and are associated with arrythmias and sudden cardiac death (Heijman et al., 2014). Calcium dynamics in cardiomyocytes were assessed using fluorescence microscopy and Fluo-4 calcium indicator. The maximum change in intracellular calcium levels did not differ significantly between FRDA cardiomyocytes and their isogenic controls (Figure 3A). However, distinct differences were observed in the regularity of calcium transients. These were categorized as either regular or irregular according to established guidelines (Gaballah et al., 2022; Hwang et al., 2020) (Figure 3B). FRDA cardiomyocytes exhibited a significantly higher incidence of irregular calcium oscillations compared to isogenic control cardiomyocytes.

**Figure 3.**
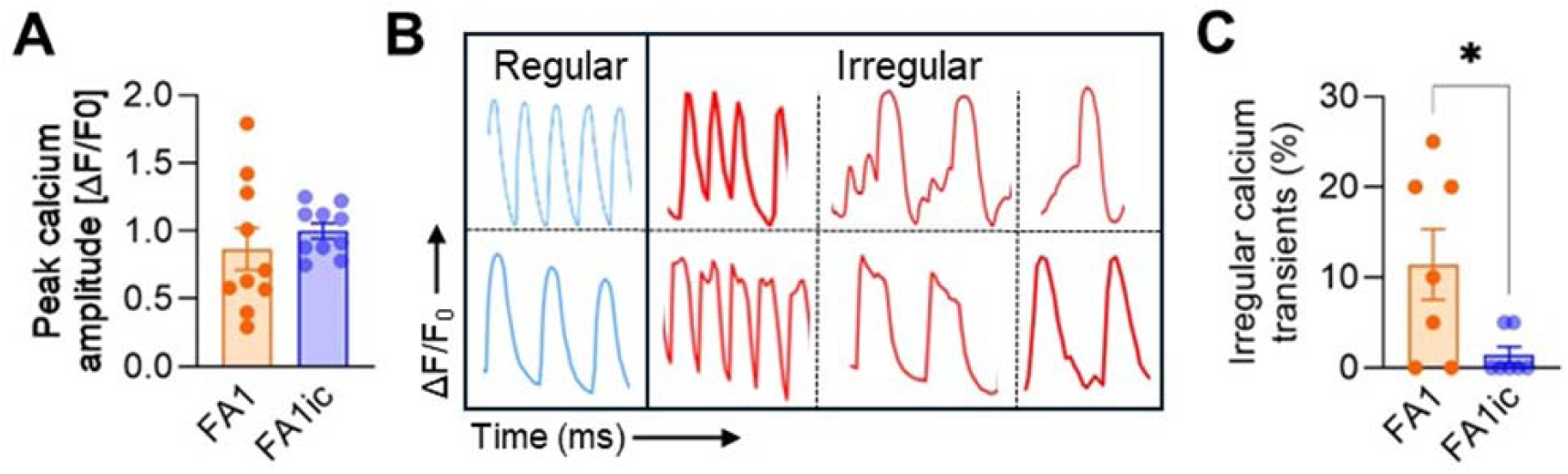
Friedreich ataxia iPSC-derived cardiomyocytes exhibit a high frequency of irregular calcium transients. (A) Peak calcium amplitude of FRDA and isogenic control iPSC-derived cardiomyocytes (n = 3-6 independent experiments). (B) Representative traces of cardiomyocyte calcium cycling. Calcium transients were categorized into either regular or irregular. (C) Percentage of cardiomyocytes exhibiting irregular calcium transients (n = 3-6 independent experiments). Data are presented as mean ± SEM. *p<0.05, **p<0.01 by unpaired t-test.

To further evaluate the arrhythmic potential of FRDA-iPSCs derived cardiomyocytes, multielectrode array recording was performed to measure the extracellular field potential of 2D cultured cardiomyocytes (Supplementary Figure 1A-B). FRDA cardiomyocytes exhibited a significantly longer field potential duration compared to isogenic controls; however, this difference was no longer significant after corrected for beating frequency (Supplementary Figure 1C-D).

### Friedreich ataxia iPSC-derived cardiomyocytes exhibit mitochondrial dysfunction

Frataxin is a mitochondrial protein essential for maintaining mitochondrial homeostasis and function, with critical implications for cardiac health. We next evaluated whether FRDA-iPSC-derived cardiomyocytes exhibited the mitochondrial dysfunction characteristic of FRDA. Cardiomyocytes derived from the FA1 iPSC line exhibited a significant increase in cell death compared to their isogenic control (FA1ic) counterparts (Figure 4A), along with an approximately 2-fold increase in mitochondrial reactive oxygen species (ROS) levels (Figure 4B). Mitochondrial membrane potential was similar between FRDA and isogenic control cardiomyocytes (Figure 4C). When mitochondrial morphology was categorized as either predominantly networked or fragmented (Figure 4D), FA1 FRDA cardiomyocytes displayed a clear shift from predominantly networked to fragmented mitochondrial morphology compared to their FA1ic isogenic controls (Figure 4E-F).

**Figure 4.**
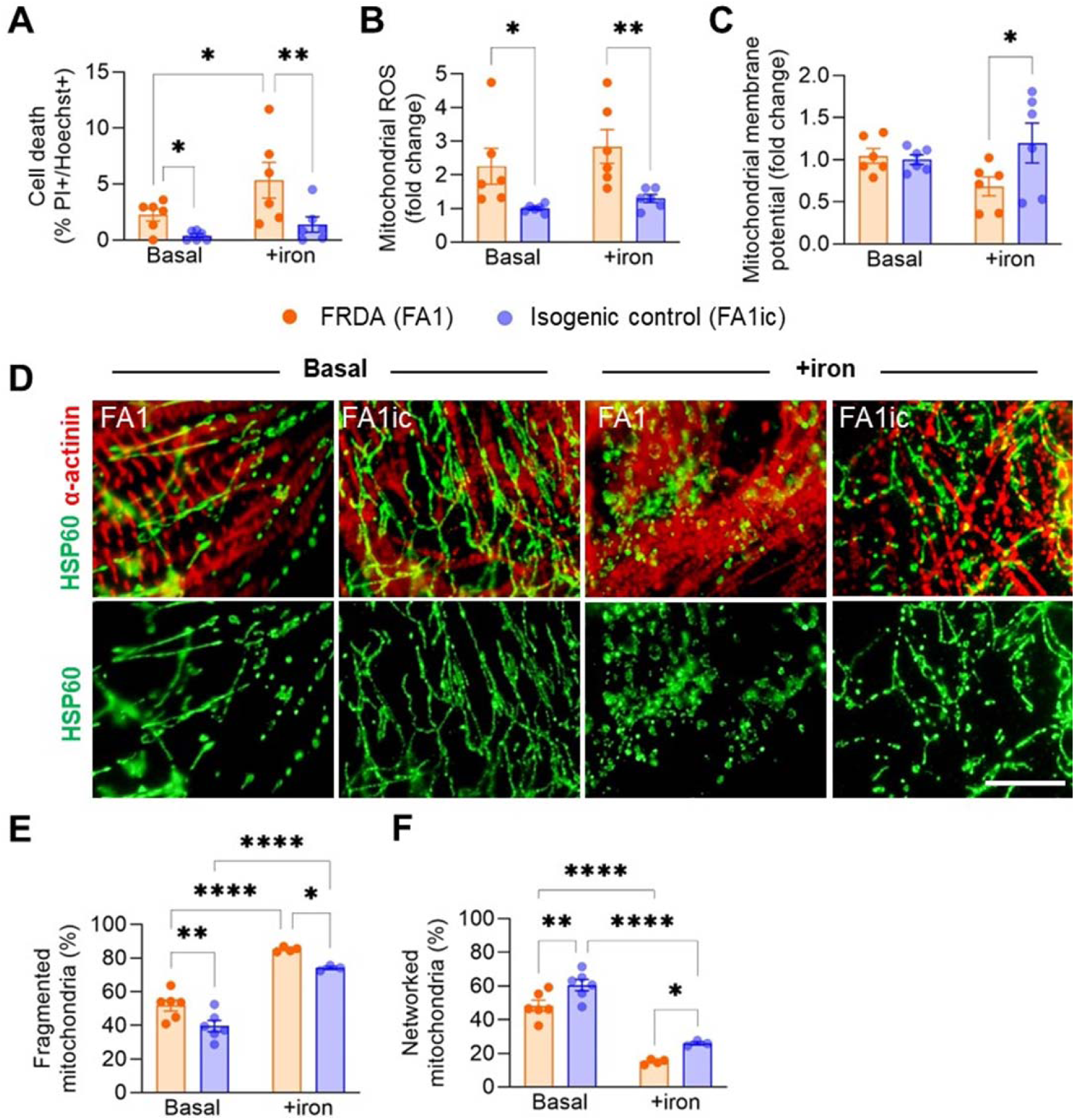
Mitochondrial dysfunction in Friedreich ataxia iPSC-derived cardiomyocytes is exacerbated by iron-induced metabolic stress. (A-F) Percentage of cell death (A), mitochondrial reactive oxygen species (ROS) levels (B), mitochondrial membrane potential (C), representative images of cardiomyocyte mitochondria stained for Hsp60 (green) and α-actinin (red) (D; scale bar = 10 µm), and quantified percentage of fragmented (E) and networked (F) mitochondrial morphologies in cardiomyocytes derived from FRDA (FA1) and isogenic control (FA1ic) iPSCs. Cells were cultured in CM medium under basal conditions or supplemented with 200 µM iron sulphate for 48 hours (+iron) (n = 3 independent experiments). Data are presented as mean ± SEM. *p<0.05, **p<0.01, ****p<0.0001 by unpaired t-test.

Deficiency in iron-sulfur cluster biogenesis, a hallmark of frataxin loss, compromises mitochondrial function and can lead to depolarization of the mitochondrial membrane potential, particularly under conditions of iron-induced metabolic stress, This phenomenon has been previously noted in mouse cerebellar granule cells (Abeti et al., 2016) and FXN-silenced human neuroblastoma cells (Bolinches-Amorós et al., 2014). To determine whether this phenotype could be recapitulated in iPSC-derived cardiomyocytes, a disease-relevant cellular stressor was applied. Elevated iron levels have been reported in the left ventricle of individuals with FDRA (Koeppen et al., 2015), and abnormal iron aggregates have been detected throughout the myocardium (Michael et al., 2006), indicating iron dysregulation.

This can be modelled *in vitro* through iron supplementation in culture (Lee et al., 2014; Lee et al., 2016). Treatment with iron sulfate for 48 hours resulted in significantly increased cell death in FRDA iPSC-derived cardiomyocytes, without notably affecting the viability of isogenic control cardiomyocytes (Figure 4A). While iron supplementation did not further elevate mitochondrial ROS levels in FRDA cardiomyocytes, which already exhibited higher ROS compared to controls (Figure 4B), it revealed a difference in mitochondrial membrane potential, with FRDA cardiomyocytes showing significantly lower mitochondrial membrane potential than their isogenic counterparts (Figure 4C). Additionally, iron-induced metabolic stress increased mitochondrial fragmentation in both FRDA and control cardiomyocytes, indicating a generalized mitochondrial stress response (Figure 4D-F).

### Friedreich ataxia iPSC-derived multicellular cardiac microtissues exhibit cellular injury and contractile abnormalities

A multicellular cardiac microtissue model, composed of cardiomyocytes along with non-cardiomyocyte populations such as endothelial cells, smooth muscle cells, autonomic neurons and cardiac fibroblasts (Figure 5A), was employed to better recapitulate the 3D microenvironment and heterogeneous cellular composition of native heart tissues. FRDA iPSC-derived cardiac microtissues showed elevated levels of lactate dehydrogenase in the conditioned media compared to isogenic control-derived microtissues, indicating increased cellular injury (Figure 5B). Consistent with findings in 2D cultures, FRDA cardiac microtissues also demonstrated contractile abnormalities (Figure 5C), including prolonged total contraction duration (Figure 5D), time to peak contraction (Figure 5E), and relaxation time (Figure 5F), as well as a reduced beating rate (Figure 5G) relative to isogenic controls. Interestingly, FRDA microtissues exhibited a higher contraction amplitude compared to controls (Figure 5H). Moreover, beat rate variability was significantly increased in FRDA iPSC-derived microtissues (Figure 5I), a feature not observed in the 2D cardiomyocyte model (Figure 2F).

**Figure 5.**
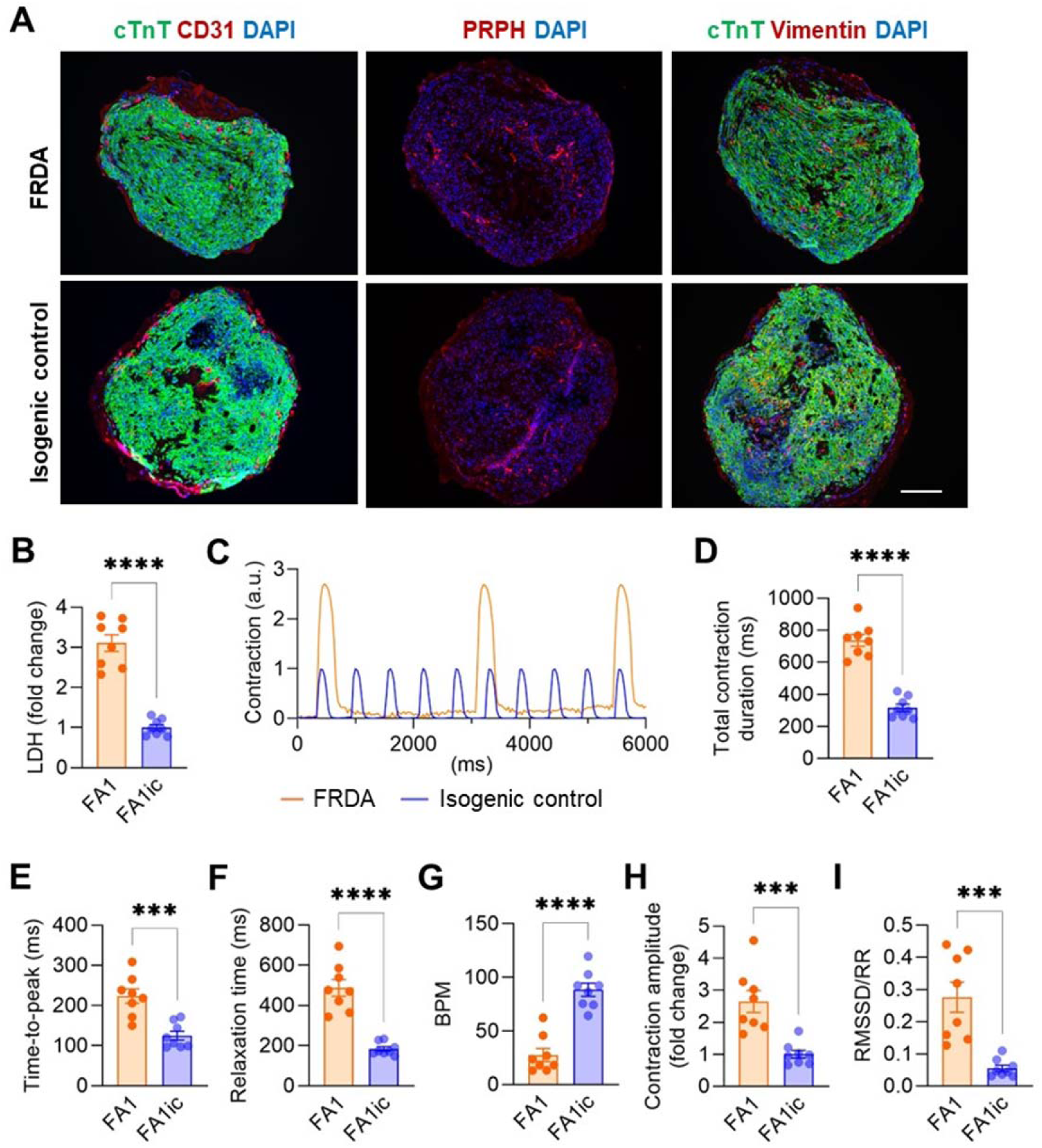
Friedreich ataxia iPSC-derived multicellular cardiac microtissues exhibit cardiac dysfunction. (A) Representative images of multicellular cardiac microtissues engineered from FRDA (FA1) or isogenic control (FA1ic) iPSCs, stained for cardiac troponin T positive cardiomyocytes (cTnT, green), CD31 positive endothelial cells (red), peripherin positive autonomic neurons (PRPH, red), and vimentin positive cardiac fibroblasts (red). Scale bar = 200 µm. (B) Lactate dehydrogenase (LDH) release in FA1 and FA1ic multicellular cardiac microtissues (n = 4 independent experiments). (C) Representative contraction trace of FA1 and FA1ic cardiac microtissues (a.u. = arbitrary units). (D-I) Quantification of contraction parameters: total contraction time (D), time to peak contraction (E), relaxation time (F), beating rate (BPM) (G), peak contraction amplitude (H), and beat rate variability (RMSSD/RR) (I) (n = 4 independent experiments). Data are presented as mean ± SEM Data are mean ± SEM. ***p<0.0001, ****p<0.0001 by unpaired t-test.

### Transcriptomic changes in Friedreich ataxia iPSC-derived cardiomyocytes

To investigate the transcriptional consequences of frataxin deficiency in cardiomyocytes, bulk RNA sequencing was performed on iPSC-derived cardiomyocytes from three FRDA iPSC lines (FA1–FA3) and their isogenic control iPSC lines (FA1ic–FA3ic) (Figure 6A-C). This approach allowed for both patient-specific comparisons and pooled analysis to identify differentially expressed genes (DEGs) associated with the FRDA cardiac phenotype. Each patient line was analyzed independently against its matched isogenic control to account for individual genetic backgrounds.

**Figure 6.**
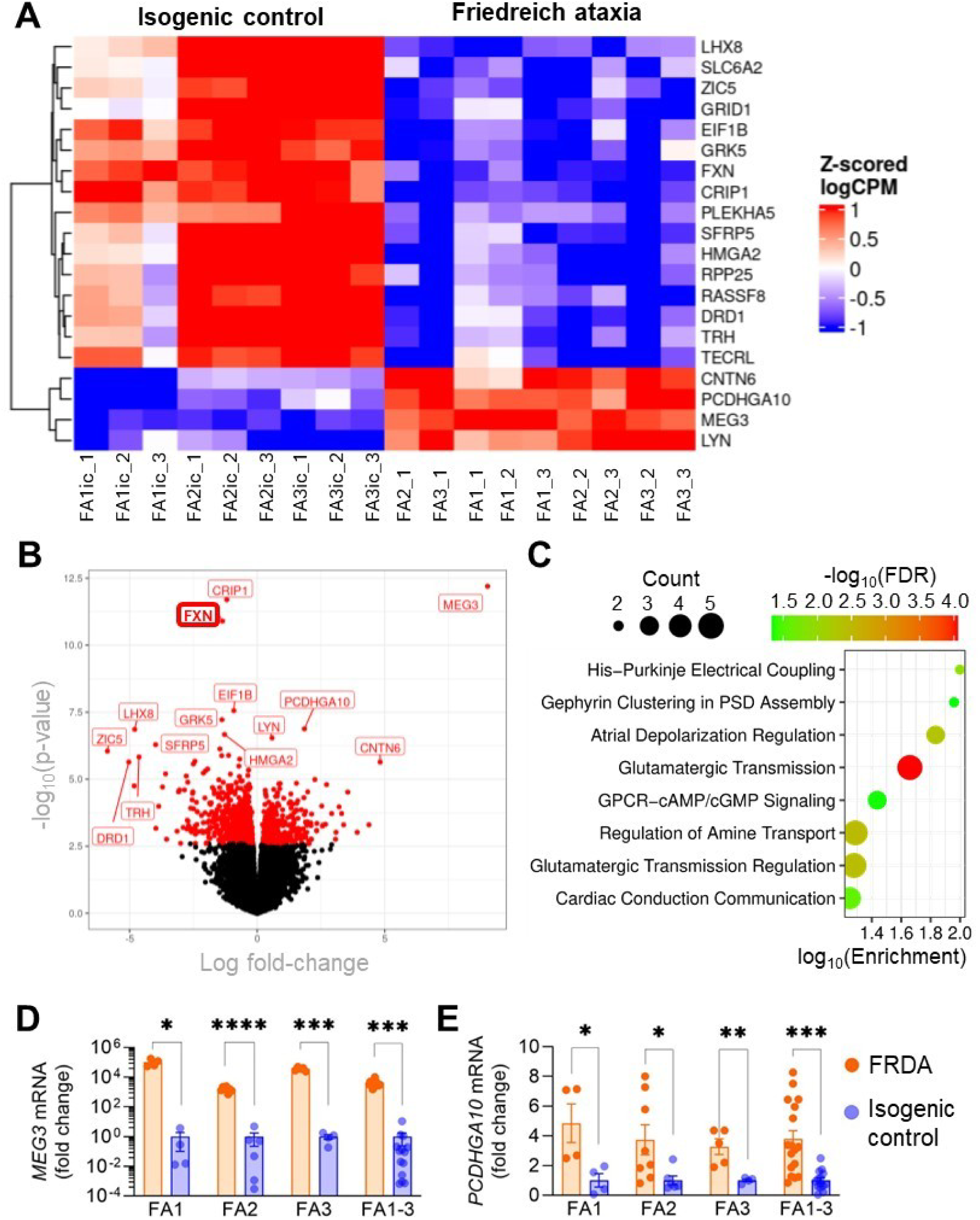
Transcriptomic analysis of cardiomyocytes derived from Friedreich ataxia and isogenic control iPSCs. (A) Heatmap of globally identified differentially expressed genes in cardiomyocytes derived from all FRDA and isogenic control iPSC lines. (B) Volcano plot showing differentially expressed genes from a combined analysis of cardiomyocytes from all three iPSC lines versus isogenic controls (FA1-3 versus FA1ic-3ic). Genes in red have an adjusted p-value (FDR) < 0.05. (C) GO Biological Processes identified using the PANTHER Overrepresentation Test, applied to differentially expressed genes with a false discovery rate < 0.05 and an absolute log fold change > 2. The top eight enriched biological processes are shown. (D-E) RT-qPCR analysis of *MEG3* (C) and *PCDHGA10* (D) mRNA expression in iPSC-derived cardiomyocytes. Data represent mean ± SEM for individual iPSC lines (FA1, FA2, FA3) and the pooled average across all 3 lines (FA1-3). *p<0.05, **p<0.01, ***p<0.001, ****p<0.0001 by unpaired t-test.

The number of DEGs varied considerably across patient lines, with 13, 2,445, and 5,280 DEGs identified in FA1, FA2, and FA3 respectively (Supplementary Tables 1-3). Of note, *MEG3* (maternally expressed gene 3) and *PCDGHA10* (protocadherin gamma subfamily A-10) were consistently dysregulated across all three FRDA patient lines, suggesting their potential as core transcriptional markers of FRDA cardiomyopathy (Figure 6A-C, Supplementary Table 4). *MEG3*, a long non-coding RNA involved in apoptosis and oxidative stress regulation, was the most significantly upregulated gene in FRDA cardiomyocytes.

*PCDHGA10*, a calcium-dependent cell adhesion molecule, was upregulated and has known roles in neuronal circuit formation and emerging relevance in cardiac tissue organization and remodeling. *MEG3* and *PCDGHA10* expression levels were further validated by qPCR (Figure 6D-E). Pooled analysis of all FRDA samples confirmed a significant downregulation of *FXN* expression in FRDA iPSC-derived cardiomyocytes (Figure 6A-B) (Table 2).

**Table 2.** Top 15 differentially expressed genes in FRDA-iPSC-derived cardiomyocytes from three individuals with FRDA, ranked by false discover rate. Negative log fold change values indicate downregulation in FRDA-iPSC-derived cardiomyocytes relative to their corrected isogenic controls.

| Gene name | Adjusted p-value | Log fold-change |
| --- | --- | --- |
| *MEG3* | 1.07e-8 | 9.03 |
| *CRIP1* | 1.78e-8 | -1.19 |
| *FXN* | 6.55e-8 | -1.36 |
| *EIF1B* | 0.000119 | -0.915 |
| *GRK5* | 0.000206 | -1.38 |
| *LHX8* | 0.000338 | -4.8 |
| *PCDHGA10* | 0.000338 | 1.85 |
| *HMGA2* | 0.000464 | -1.28 |
| *LYN* | 0.000551 | 0.584 |
| *SFRP5* | 0.000882 | -3.98 |
| *TECRL* | 0.00115 | -1.46 |
| *ZIC5* | 0.00129 | -5.87 |
| *RASSF8* | 0.00163 | -1.39 |
| *RPP25* | 0.00163 | -1.12 |
| *TRH* | 0.0017 | -4.63 |

Additional DEGs were identified, many of which are implicated in calcium dysregulation (e.g. *TECRL*, *TRH*, *SLC6A2*, *GRID1*), contractile and electrophysiological dysfunction (e.g. *ERK5*, *LYN*, *LHX8*, *PCDHGA10*, *CNTN6*, *PLEKHA5*, *RASSF8*), mitochondrial dysfunction and oxidative stress (e.g. *MEG3*, *EiF1B*, *HMGA2*, *ZIC5*), and cellular stress and apoptosis (e.g. *CRIP1*, *RPP25*, *SFRP5*, *HMGA2*, *ZIC5*) (Table 2), consistent with enrichment of GO biological processes related to synaptic signaling, cardiac conduction, and ion transport (Figure 6C; Supplementary Table 5). Collectively, these transcriptomic changes provide insight into the potential molecular mechanisms underlying FRDA cardiomyopathy and identify candidate genes for further functional studies and potential therapeutic targeting.

## Discussion

This study presents the development and characterization of a human iPSC-based model of FRDA cardiomyopathy, using three patient-derived iPSC lines and their CRISPR/Cas9-corrected isogenic controls. Restoration of *FXN* expression through targeted genome editing enabled the isolation of the effects of frataxin deficiency on cardiomyocyte function and gene regulation. The model revealed early and intrinsic cardiac abnormalities in FRDA, including contractile dysfunction, impaired calcium handling, compromised viability, mitochondrial oxidative stress, mitochondrial fission, and transcriptional dysregulation. Mitochondrial disease phenotypes were further exacerbated under iron-induced metabolic stress conditions, highlighting the metabolic vulnerability of FRDA cardiomyocytes. Transcriptomic analysis identified a distinct molecular signature, with *MEG3* and *PCDHGA10* consistently dysregulated across all FRDA lines. Together, these results provide mechanistic insight into the early stages of FRDA cardiomyopathy and highlight potential novel targets for therapeutic intervention.

Cardiac manifestations are a primary cause of mortality in FRDA, with structural remodeling and arrhythmias often evident early in disease progression (Weidemann et al., 2013). Despite this, the underlying cellular mechanisms remain incompletely understood (Payne, 2022).

Previous studies using FRDA iPSC-derived cardiomyocytes have reported abnormalities in contractility and calcium handling (Crombie et al., 2017; Wong et al., 2019); however, the absence of isogenic controls in those studies limited the ability to attribute observed phenotypes specifically to *FXN* deficiency. *In vitro* modeling of FRDA is challenged by inter-patient genetic variability and phenotypic heterogeneity. This limitation was addressed in the present study through CRISPR/Cas9-mediated excision of the pathogenic GAA repeat expansion in the *FXN* gene, which restored frataxin mRNA and protein levels and reversed multiple contractile and mitochondrial abnormalities. Although this correction yields a genetic state not typically found in healthy individuals, who usually carry 8 to 30 GAA repeats (Pandolfo, 2008), the resulting isogenic control iPSCs expressed frataxin at levels comparable to healthy donor-derived iPSCs, supporting their validity as physiologically relevant controls.

The availability of genetically matched control lines enables precise characterization of FRDA-associated cardiac phenotypes. Evidence of contractile dysfunction, including prolonged contraction and relaxation times, across multiple FRDA lines indicates that both systolic and diastolic impairments are intrinsic features of frataxin deficiency.

The prolonged time-to-peak contraction observed in our FRDA cardiomyocytes further aligns with similar findings in failing human myocardium (Davies et al., 1995), reinforcing the pathological relevance of our model and its utility for studying early-stage FRDA cardiomyopathy. In our 3D cardiac microtissue model, FRDA tissues exhibited both prolonged contraction duration and increased contraction amplitude, despite a lower beat rate. This pattern contrasts with the typical force-frequency relationship observed in adult cardiomyocytes, where lower beating rates are generally associated with a reduced contraction amplitude (Davies et al., 1995). The observed discrepancy may reflect the immaturity of iPSC-derived cardiomyocytes and the absence of external pacing and physiological loading *in vitro*. Additionally, the lack of neurohumoral regulation in our system such as autonomic input and circulating factors may contribute to differences in contraction behavior compared to *in vivo* conditions, where systemic influences modulate cardiac function.

Heart rate itself is a notable variable in FRDA pathology, particularly when interpreting systolic function (Peverill et al., 2017). Although our FRDA models consistently showed lower BPM, clinical studies report elevated heart rates in individuals with FRDA with preserved left ventricular ejection fraction (Dedobbeleer et al., 2012; Mottram et al., 2011). Despite this elevated heart rate, individuals with FRDA exhibit reduced systolic and early diastolic peak velocities, suggesting intrinsic myocardial dysfunction. Interestingly, frataxin knockout mice show reduced heart rate and contraction amplitude, with partial normalization of heart rate following frataxin restoration, but persistent contractile impairment (Vyas et al., 2012). This suggests that frataxin deficiency induces lasting intrinsic defects in cardiac contractility that may not be fully reversible, even when frataxin levels are restored. These findings highlight species-specific differences and underscore the importance of considering both cell-autonomous and systemic regulatory mechanisms when interpreting cardiac phenotypes in FRDA.

Aberrant calcium handling emerged as a key pathological feature in our FRDA cardiomyocytes. Although calcium transient amplitude remained unchanged and beat rate variability was not elevated in 2D monolayer cultures, an increased incidence of irregular calcium oscillations was observed, indicating disrupted excitation–contraction coupling in FRDA iPSC-derived cardiomyocytes. Interestingly, increased beat rate variability was detected in 3D multicellular cardiac microtissues derived from the same FRDA lines, suggesting that arrhythmic risk may be amplified in the presence of a more physiologically complex tissue environment. This finding raises the possibility that non-cardiomyocyte populations, such as cardiac fibroblasts and autonomic neurons, may contribute to the emergence of electrical instability in FRDA cardiac tissue. The observed irregular calcium behavior may reflect early arrhythmogenic potential, aligning with clinical reports of conduction abnormalities and atrial arrhythmia in individuals with FRDA (Payne, 2022). Mechanistically, frataxin deficiency has been linked to impaired mitochondrial calcium uptake and ATP depletion, potentially compromising the activity of calcium-handling proteins such as SERCA and the sodium/calcium exchanger (Bolinches-Amorós et al., 2014; Crombie et al., 2017). This dysfunction may lead to cytosolic sodium accumulation, reversal of exchanger activity, and increased ROS, which further destabilize calcium channels and promote arrhythmogenesis (Li et al., 2014). Downregulation of solute carrier genes encoding calcium exchangers in FRDA fibroblasts (Napierala et al., 2017) supports the notion of systemic calcium dysregulation in FRDA.

Frataxin, a mitochondrial protein essential for iron sulphur cluster biogenesis and mitochondrial homeostasis, is known to regulate oxidative metabolism and redox balance (Santos et al., 2010). Its deficiency in FRDA has been shown to impair mitochondrial function, elevate ROS levels, and reduce mitochondrial membrane potential across various models, including yeast, animal models, and human cell lines (Akbari et al., 2019; Campuzano et al., 1996; Radisky et al., 1999). In the current model, FRDA iPSC-derived cardiomyocytes exhibited increased mitochondrial ROS levels, elevated cell death, and a shift toward fragmented mitochondrial morphology, consistent with the established role of frataxin in maintaining redox balance and mitochondrial integrity (Bolinches-Amorós et al., 2014; Hanson et al., 2019). Despite these abnormalities, mitochondrial membrane potential remained unchanged under baseline conditions, a finding in agreement with previous reports in primary fibroblasts isolated from individuals with FRDA (Abeti et al., 2018). This may reflect compensatory mechanisms active during early cardiomyocyte development or the presence of a threshold that must be surpassed to trigger depolarization. To test this, iron overload was applied, a disease-relevant stressor that mimics metabolic stress due to myocardial iron accumulation observed in FRDA (Koeppen et al., 2015; Michael et al., 2006). Under these conditions, mitochondrial membrane depolarization, cell death and mitochondrial fragmentation were significantly exacerbated, with FRDA cardiomyocytes consistently exhibiting more severe phenotypes. These findings suggest that frataxin-deficient cardiomyocytes are predisposed to mitochondrial dysfunction, with iron dysregulation acting as a catalyst for disease progression.

In addition to mitochondrial stress, evidence of early pathological remodeling was observed, including cardiomyocyte hypertrophy and increased apoptosis. Cardiac structural remodeling is a hallmark of FRDA cardiomyopathy and has been reported in both iPSC-derived cardiomyocytes and transgenic mouse models (Hick et al., 2013; Puccio et al., 2001). The increased cardiomyocyte size observed in the present study reflects this hypertrophic phenotype and aligns with clinical observations of ventricular wall thickening in individuals with FRDA, although it represents only one component of the broader structural remodeling seen *in vivo* (Peverill et al., 2019). The concurrent increase in cell death further aligns with autopsy findings of cardiomyocyte loss in FRDA hearts (Koeppen et al., 2015), supporting a model in which cumulative cell loss contribute to progressive cardiac remodeling and contractile dysfunction (Schreiber et al., 2019).

To further elucidate the molecular basis of these phenotypes observed in FRDA iPSC-derived cardiomyocytes, RNA sequencing was performed on cardiomyocytes derived from three individuals with FRDA and their isogenic controls. This approach enabled both individual and pooled analyses of DEGs, revealing broad transcriptional changes associated with frataxin deficiency. Pooled analysis identified dysregulation of genes involved in calcium signalling, cardiac rhythm, and hypertrophy, including *CRIP1* (calcium binding and cytoskeletal organization) (Ye et al., 2025), *GRK5* (β-adrenergic signalling and hypertrophy) (Pfleger et al., 2019), and *TECRL* (arrhythmia and calcium handling) (Porretta et al., 2025).

GO enrichment analysis revealed significant overrepresentation of biological processes related to cardiac conduction, synaptic signalling, and membrane depolarization, processes essential for coordinated electrophysiological activity and neuro-cardiac integration. These changes align with the functional abnormalities observed in FRDA cardiomyocytes and underscore the widespread impact of reduced frataxin.

Among all DEGs, *MEG3* and *PCDHGA10*, were consistently dysregulated across all three FRDA iPSC lines, suggesting they may represent core components of the FRDA cardiac transcriptional signature. *MEG3* (maternally expressed gene 3) emerged as the most significantly upregulated transcript. This long non-coding RNA is known to regulate diverse cellular processes, including apoptosis, angiogenesis, and ferroptosis (Huang & Chen, 2012; Maimaitizunong et al., 2022). Elevated expression of *MEG3* in FRDA cardiomyocytes may contribute to increased cell death and oxidative stress. In cardiovascular contexts, *MEG3* downregulation has been shown to reduce cardiomyocyte apoptosis following myocardial infarction (Wu et al., 2018) and reduce fibrosis in cardiac fibroblasts (Li et al., 2021), indicating a potential pro-apoptotic and pro-fibrotic role in cardiac tissue. Therefore, upregulation of *MEG3* in FRDA may represent a maladaptive stress response that exacerbates mitochondrial dysfunction and cell loss, highlighting its potential role in the pathogenesis of FRDA cardiomyopathy. Interestingly, *MEG3* represents the second long non-coding RNA implicated in FRDA, following identification of the long non-coding RNA *TUG1* as a potential blood-based early biomarker (Koka et al., 2024). While *TUG1* was significantly downregulated in FRDA patient blood and serum, MEG3 showed robust upregulation in cardiomyocytes, suggesting distinct roles in FRDA pathogenesis and the potential for complementary biomarker development.

*PCDHGA10* (protocadherin gamma subfamily A-10), was also significantly upregulated in FRDA cardiomyocytes. *PCDHGA10* is a calcium-dependent cell adhesion molecule implicated in the formation and maintenance of neuronal circuits (Liu et al., 2009).

Dysregulation of *PCDHGA10* has been linked to neurodegeneration in Huntington’s disease (An et al., 2012) and proprioceptive neuron degeneration in FRDA (Dionisi et al., 2023). A recent transcriptomic study has also suggested a role for *PCDHGA10* in cardiac development and pathology, with increased expression in cardiomyocytes correlating with disrupted muscle fibre organization and contraction (Lu et al., 2023). Its upregulation in FRDA cardiomyocytes may therefore contribute to impaired structural organization and intracellular communication. Collectively, the dysregulation of *MEG3* and *PCDHGA10*, genes implicated in oxidative stress, apoptosis, disrupted calcium signaling, and structural remodeling, reflects key pathological processes underlying FRDA cardiomyopathy. Their consistent expression patterns across patient lines highlight their potential as molecular markers of FRDA cardiomyopathy and underscore the value of transcriptomic profiling in identifying targets for mechanistic and therapeutic investigation.

### Conclusion

This study provides a comprehensive characterization of cardiomyocyte pathology in FRDA using patient-derived iPSC models alongside CRISPR/cas9-corrected isogenic controls. Frataxin deficiency was found to cause early and intrinsic cardiomyocyte dysfunction, characterized by impaired contractility, irregular calcium handling, elevated mitochondrial oxidative stress, mitochondrial fission, and increased cell death. These pathogenic phenotypes were evident under baseline conditions and were further exacerbated by iron-induced metabolic stress conditions. Transcriptomic profiling revealed a distinct molecular signature associated with *FXN* downregulation, with *MEG3* and *PCDHGA10* emerging as consistently dysregulated genes across all patient lines. The use of genetically matched isogenic iPSC models enabled precise dissection of disease-specific phenotypes and highlights their utility as a platform for uncovering pathogenic mechanisms and guiding the development of targeted therapeutic strategies.

## Statements and Declarations

### Data Availability

The Gene Expression Omnibus accession number for the RNA-seq dataset reported in this paper is GSE305638.

### Author contributions

J.G.L. and S.Y.L. conceived the project. J.G.L., A.P., M.D., L.C., M.D., R.P., M.N., and S.Y.L. were responsible for funding acquisition and experimental design. J.G.L., H.Z., L.J., A.M.K., R.J.P., L.L., N.S., A.S.M., S.W., J.S.N., and S.Y.L. performed experiments and data curation. J.G.L., H.Z., L.J., A.M.K., R.J. P., L.L., S.W., J.M.P., and S.Y.L. analyzed data. J.G.L., H.Z., L.J., L.L., A.P., M.D., L.C., M.D., R.P., D.M., M.N., and S.Y.L. interpreted results. J.G.L., H.Z., L.J., L.L., J.M.P., and S.Y.L. prepared figures. J.G.L. and S.Y.L. drafted manuscript. All authors participated in manuscript revision and approved the final version of the manuscript.

