## Supplementary Materials for "Frataxin deficiency drives cardiac dysfunction and transcriptional dysregulation in Friedreich ataxia iPSC model"

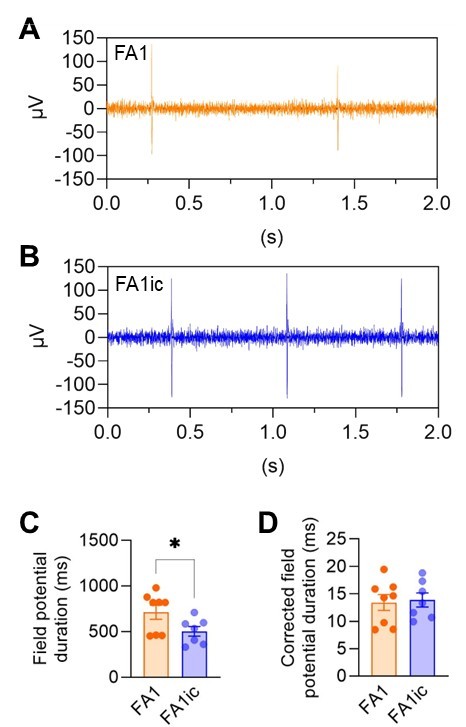

**Supplementary Figure 1. Extracellular field potential of Friedreich ataxia iPSC-derived cardiomyocytes.** (A-B) Representative trace of extracellular field potential of FA1 (A) and FA1ic (B) cardiomyocytes recorded from multielectrode array. (C-D) Field potential duration (C) and corrected field potential duration (D) of FA1 and FA1ic cardiomyocytes (n = 4 independent experiments).

**Supplementary Table Legends**

**Supplementary Table 1.** Differential expression analysis results for cardiomyocytes from patient 1 (FA1), generated using edgeR within the ARMOR pipeline.

**Supplementary Table 2.** Differential expression analysis results for cardiomyocytes from patient 2 (FA2), generated using edgeR within the ARMOR pipeline.

**Supplementary Table 3.** Differential expression analysis results for cardiomyocytes from patient 3 (FA3), generated using edgeR within the ARMOR pipeline.

**Supplementary Table 4.** Differential expression analysis results from the combined analysis of cardiomyocytes from patients 1–3 (FA1–FA3), generated using edgeR within the ARMOR pipeline.

**Supplementary Table 5.** GO Biological Process enrichment from PANTHER Overrepresentation Test on combined patient cardiomyocyte data (FDR < 0.05, |Fold Change| ≥ 2).
